# *C11orf70* mutations causing primary ciliary dyskinesia disrupt a conserved step in the intraflagellar transport-dependent assembly of multiple axonemal dyneins

**DOI:** 10.1101/211953

**Authors:** Mahmoud R. Fassad, Amelia Shoemark, Pierrick le Borgne, France Koll, Mitali Patel, Mellisa Dixon, Jane Hayward, Charlotte Richardson, Emily Frost, Lucy Jenkins, Thomas Cullup, Eddie MK Chung, Michel Lemullois, Anne Aubusson-Fleury, Claire Hogg, David R. Mitchell, Anne-Marie Tassin, Hannah M. Mitchison

## Abstract

Primary ciliary dyskinesia (PCD) is a genetically and phenotypically heterogeneous disorder characterized by destructive respiratory disease and laterality abnormalities due to randomised left-right body asymmetry. PCD is mostly caused by mutations affecting components of the core axoneme structure of motile cilia that are essential for cilia movement. In addition, there is a growing group of PCD genes that encode proteins essential for the assembly of the ciliary dynein motors and the active transport process that delivers them from their cytoplasmic assembly site into the axoneme. We screened a cohort of affected individuals for disease-causing mutations using a targeted next generation sequencing panel and identified 2 unrelated families (3 affected children) with mutations in the uncharacterized *C11orf70* gene. The affected children share a consistent PCD phenotype from early life with laterality defects and immotile respiratory cilia displaying combined loss of inner and outer dynein arms (IDA+ODA). Phylogenetic analysis shows *C11orf70* is highly conserved, distributed across species similarly to proteins involved in the intraflagellar transport (IFT)-dependant assembly of axonemal dyneins. *Paramecium C11orf70* RNAi knockdown led to combined loss of ciliary IDA+ODA with reduced cilia beating and swim velocity. Fluorescently tagged C11orf70 in *Paramecium* and *Chlamydomonas* localises mainly in the cytoplasm with a small amount in the ciliary component, its abundance in the axoneme being IFT-dependant. During ciliogenesis, C11orf70 accumulates at the ciliary tips in a similar distribution to the IFT-B protein IFT46. In summary, C11orf70 is essential for IFT-dependant assembly of dynein arms and *C11orf70* mutations cause defective cilia motility and PCD.

## Introduction

Cilia are complex organelles that form a distinct cellular compartment projecting from the surface of non-dividing cells. They have a microtubular-based axoneme core structure, contained within a specialised membrane that extends from the plasma membrane. This basic structure is highly conserved across species to serve various physiological functions.^1^ Motile and non-motile types of the cilia can be distinguished in the human body that have differences in their axoneme structural arrangement and motilities.^2; 3^ Non-motile primary cilia present on most cells contain nine pairs of microtubules forming a circular scaffold of nine-fold symmetry termed 9+0. Nodal cilia of the embryonic left-right organiser (node) have a similar structure but with dynein ‘arm’ motor subunits attached to the peripheral microtubules that provide motility machinery for ciliary beating.^4^ Motile cilia of the respiratory epithelium, fallopian tubes and the brain ependyma which lines the ventricular system have a 9+2 arrangement, where the peripheral microtubules attached to dynein arm motors surround a central pair of microtubules. The sperm flagellum has the same structure as motile cilia with subtle differences in the distribution of dynein arms along the axoneme.^5^ Lastly, kinocilia of the inner ear hair cells have a 9+2 organization but lack beating motility, containing outer but not inner dynein arms.

The main function of cilia/flagella motility is in generating fluid flow, or movement within fluids. Defective cilia motility leads to primary ciliary dyskinesia (PCD, MIM: 244400) a genetic disorder in which left-right positioning of body organs is randomised such that laterality problems affect about half of affected individuals. Such laterality defects may be associated with -in some cases-cardiac, spleen and other organ problems.^6^ Children with PCD suffer from chronic, widespread upper and lower airway infections and congestion, progressing towards bronchiectasis. Symptoms often present early in life with neonatal respiratory distress syndrome. Affected individuals can have hydrocephalus, hearing defects, retinal dystrophy, and infertility in adults especially in males due to sperm dysmotility.^7; 8^

PCD is inherited as an autosomal recessive or X-linked disease. The genetic cause of PCD, where known, has mainly been mutations affecting in one of the hundreds of genes encoding essential components of the complex motile ciliary machinery. PCD is dominated by mutations affecting the axonemal dynein arm motors, for example in subunits of the outer dynein arm (ODA) such as *DNAH5*,^9^ or in proteins involved in ODA targeting and anchoring such as *CCDC103*, *ARMC4* and ODA docking complex (ODA-DC) components such as *CCDC151*.^10–16^.

Perhaps more surprising than the finding that mutations in cilia structural components are a frequent cause of PCD has been a growing number of PCD loci that do not encode parts of the cilium per se, but rather encode proteins that act in the apparently complicated process of the assembly and transport of ciliary dynein motors.^17^ Many such assembly factors work strictly in the cytoplasm in a chaperoning step and cause PCD when mutated, such as *DNAAF1*,^18–20^ *DNAAF2*,^21^ *DNAAF3*,^22^ *DYX1C1* (*DNAAF4*),^23; 24^ *HEATR2* (*DNAAF5*),^25; 26^ *LRRC6*^27–29^ and *PIH1D3*.^30; 31^ Other factors, identified in both the human population through analysis of PCD patients and in genetic screens for cilia motility defects in model organisms, resemble chaperones in that they affect the assembly of multiple ciliary dynein isoforms, but also resemble transport factors based on their presence in a detergent-soluble ciliary compartment. These include *ZMYND10*^32; 33^ and *C21orf59*.^34; 35^

A number of additional proteins involved in dynein assembly have been identified from non-human studies including *ODA8*, *ODA5* and *ODA10* of the flagellated model organism *Chlamydomonas reinhardtii*. *ODA5* and *ODA10* directly interact as a complex required for formation of an assembly– competent outer dynein arm in the cytoplasm, participating along with *ODA8* in the later stages of cytoplasmic ODA assembly.^36; 37^ ODAs are then trafficked into cilia for assembly onto the ODA-DC using the internal cargo transport system, intraflagellar transport (IFT) and subsequent steps involve proteins thought to be involved in an IFT-dependent trafficking step, such as ODA16 (*DAW1* or *WDR69* in human) which has so far not been linked to disease in humans.^38–40^ The anterograde or base-tip direction of cilia cargo transport involves members of the IFT-B subcomplex, one of which (IFT46) interacts with ODA16. ODA16 therefore seems to function as an adaptor between IFT-B and the ODA, required for the IFT-based transport of ODAs into the cilia/flagella compartment.^39^

To reach a genetic diagnosis of PCD, bi-allelic mutations in autosomal recessive or hemizygous state in an X-linked gene should be identified. Reported mutations in known genes account for only about 70% of PCD patients, so additional genes are still to be identified.^17^ Here we describe a novel PCD gene, *C11orf70*, and demonstrate that it encodes a protein needed for assembly of both outer and inner arm axonemal dyneins in respiratory cilia, a hallmark of the axonemal dynein assembly co-chaperones. Using model organisms, we show that *C11orf70* orthologs perform a conserved dynein assembly function and are present in both the cytoplasm and in cilia. Furthermore, ciliary abundance of this protein is detergent-sensitive and dependent on continued IFT trafficking, as expected for a factor important for the late steps in ciliary dynein assembly.

## Materials and Methods

### Subjects and families

The study was ethically approved by the London Bloomsbury Research Ethics Committee (08/H0713/82). Written informed consent was obtained from all participants or their guardians prior to enrolment in the study. Patients were diagnosed with PCD through standard diagnostic screening by the Royal Brompton Hospital PCD diagnostic service, according to European Respiratory Society diagnostic guidelines.^41^

### Next generation sequencing

Genomic DNA was extracted from whole blood samples using a standard phenol-chloroform protocol. Genetic screening was done as previously described using a targeted next generation sequencing panel including all known PCD genes, known genes causing isolated heterotaxy and other candidate genes that predicted to play a role in cilia motility.^30^ The panel was designed using the Agilent SureDesign tool (Agilent Technologies, CA, USA) to capture all coding regions of the included genes and 25 bp at the exon-intron boundaries. Library preparation was done using the SureSelectQXT Target Enrichment Kit (Agilent Technologies). Paired end sequencing (2 × 150bp) was performed using either NextSeq or HiSeq platforms (Illumina, Inc., CA, USA). A multiplex of 48 patient samples were sequenced per flow cell in each run using barcoding indices. Sequencing data were subjected to quality control and further analysis using an in-house pipeline as previously described.^42^ Alignment of Fastq sequence files with the human genome reference (GRCh37/hg19) used BWA (0.6.1-r104),^43^ with variant calling performed using Freebayes^44^ and variant annotation using Alamut^®^ Batch (Interactive Biosoftware, Llc., Rouen, France) softwares. VCF files were then converted into Excel format to enable manual variant filtration and prioritization. Only coding nonsynonymous and splicing variants (±5bp from exon-intron boundary) were retained for further analysis. Common sequence variants were then filtered out based on their minor allele frequency (MAF) to exclude all variants with MAF ≥ 0.01 in the dbSNP build 141, ExAC, Exome Variant Server and 1000 Genomes databases. Retained variants were evaluated and prioritized based on their predicted impact at the protein level and predicted pathogenicity scores using a number of in silico softwares: Human Splicing Finder, SIFT, Polyphen-2, Mutation Taster, Combined Annotation Dependent Depletion (CADD) score and Multivariate Analysis of Protein Polymorphism (MAPP). Conservation across species was studied using PhastCons, PhyloP and the BLOSUM62 matrix within BLAST.

### Sanger sequencing

Prioritized variants in *C11orf70* were further confirmed in the probands and segregated within other family members by Sanger sequencing. Primers flanking the exon harbouring the mutations were designed using the NCBI Primer-BLAST tool with genomic sequence amplification performed using standard PCR conditions and annealing temperatures based on the primer Tm temperature. Primers are listed in **Table S1**. Analysis of the sequencing data used SnapGene software (GSL Biotech LLC, Chicago, USA).

### Air-liquid interface (ALI) culture

BMI1 transformed normal human bronchial epithelial cells (BMI1 NHBE) were cultured at air liquid interface on collagen-coated transwell inserts (passage 7) for 4 weeks as previously described.^45; 46^

### Quantitative RT-PCR

Total RNA was extracted using Trizol Reagent (Qiagen Inc., Hilden, Germany). cDNA was produced using Omniscript RT kit (Qiagen Inc., Hilden, Germany). Real-time qPCR reactions were done in triplicate using the CFX96 touch real-time PCR system and iQ™ SYBR green Supermix and data analysis using Bio-Rad CFX Manager 3.1 software (Bio-Rad Labs Inc., CA, USA). Primers used for qPCR in human and *Paramecium* cells are listed in **Table S1**.

### Transmission Electron Microscopy (TEM) and immunofluorescence in human subjects

Nasal brushings from patients were fixed in 4% glutaraldehyde for transmission electron microscopy. Processing and quantification was as previously described.^47^ Briefly, samples were bound in agar, post fixed in osmium tetroxide, dehydrated, and embedded in araldite before being cut into thin sections for analysis. Sections were stained with 2% methanoic uranyl acetate and lead citrate to provide contrast and assessed on a Jeol 1400+ transmission electron microscope. Images of cilia cross sections were acquired with a digital camera (AMT 16X, Deben UK). Immunofluorescence for PCD diagnosis was conducted as previously described.^48^ All antibodies used in this study are listed in **Table S2 and S3**.

### High Speed Video Microscopy analysis (HSVM)

Nasal epithelial cells were re-suspended in Medium 199 (pH7.2, Sigma-Aldrich Co. Ltd., Dorset, UK) and assessed by light microscopy at 37°C in a 1 mm chamber. Well ciliated continuous epithelial strips were selected for assessment by high speed video. Cilia were recorded using a 100X oil immersion lens on an upright DM60 Leica microscope (Leica Microsystems GmbH, Wetzlar, Germany) at 500 frames per second (Troubleshooter, Fastec Imaging, CA, USA). Ciliary beat pattern was assessed and ciliary beat frequency calculated according to the number of frames required to complete 10 full beats.^49^

### *Paramecium* gene silencing

We identified the *Paramecium C11orf70* ortholog as *GSPATG00011350001* using public databases CILDB and ParameciumDB. A 561 bp long DNA segment corresponding to the N-terminus of the gene was cloned in L4440 plasmid between the two convergent T7 promotors.^50^ Possible RNAi off-target effects were assessed using the ParameciumDB RNAi off-target tool. *E. coli* HT115 bacteria (lacking RNAase III activity) were transformed with this silencing vector and grown in Lysogeny broth (LB) medium. Paramecia (*Paramecium tetraurelia d4-2* strain) were fed with the transformed bacteria over 3 days as previously described.^51^ The RNAi experiment was repeated several times to ensure reproducibility of the results (n>10). RNAi of the *ND7* gene, which affects trichocyst exocytosis without affecting cilia motility, was used as a negative control. Knockdown of *ND7* was attested by trichocyst retention after picric acid treatment. ^52^ To assess the degree of gene knockdown at the RNA level, a mass culture protocol was used to get sufficient quantities of paramecia for RNA extraction as previously described ^53^. Total RNA was extracted using the RNAeasy Micro kit (Qiagen). cDNA was produced using 1 μg RNA, Superscript IIII (Invitrogen) and random hexanucleotides (Invitrogen). qPCR was performed to assess the degree of knockdown at the RNA level (primers listed in **Table S1**).

### *Paramecium* cilia function tests

*Paramecium* velocity was tested using 3-5 paramecia per test transferred into a drop of conditioned BHB solution and tracked for 10 seconds every 0.3 second to assess the swimming pattern under dark field microscopy, using MetaVue software. ImageJ software was used for image analysis and measurement of the swimming velocity.^54^ For *Paramecium* cilia beating analysis, knockdown cells were adhered to a slide covered with Cell-TakTM (BD Bioscience, San Jose, CA) according to Bell et al 2015 after silencing for 72 hr as described.^55^ Cilia beating was recorded using a 63X oil immersion lens on an upright DM60 Leica microscope the same as for the nasal epithelia cells, at 500 frames per seconds at 37°C. The CiliaFA plugin in ImageJ was used for assessment of cilia beat frequency in *Paramecium*.^56^

### *Paramecium* TEM and immunofluorescence analysis

The sample fixation and processing protocol for TEM in *Paramecium* was as previously described.^57^ Images of cilia cross sections were acquired with a digital camera (AMT 16X, Deben UK). The numbers of the missing IDA and ODA arms were recorded per cross section by an observer blinded to the condition. For immunofluorescence studies, paramecia were fixed in 2% paraformaldehyde in PHEM buffer (Pipes 60mM, Hepes 25 mM, EGTA 10mM, MgCl2 2mM, adjusted to pH 6.9 with NaOH) for 15 minutes. After fixation, cells were permeabilised for 15 minutes in 1% Triton X-100 in PHEM. Cells were washed 3 times in PBS/BSA 3%. Immunostaining using polyglutamylated tubulin antibodies (1/500) as previously described.^58^

### Phylogenetic comparisons

The presence or absence of *C11orf70* orthologs was determined by searches of selected organism genomes (at the genus level) using BLASTp at NCBI with the *Chlamydomonas* ortholog protein *FBB5* sequence (XP_001694050) as query. Apparent hits were confirmed only if they also gave *FBB5* as the top hit in a reciprocal BLAST search. This procedure was also used to establish the presence or absence of two ODA subunits (HCgamma and IC2), two IDA subunits (I1HCalpha and IC140), two IFTA subunits (*IFT122* and *IFT140*), four IFTB subunits (*IFT46*, *IFT52*, *IFT88* and *IFT172*), ODA-DC2 and *ODA16*. Retention of additional IFT subunits by selected organisms was taken from van Dam et al. 2013.^59^

### Functional characterizations of *C11orf70* in *Paramecium*

The *Paramecium C11orf70* ortholog (*GSPATT00011350001*) was cloned upstream of GFP into the *Spe1-Xho1* sites of pPXV-GFP plasmid which contains the constitutive regulators of the *Paramecium* calmodulin gene.^60^ Three glycine codons were added between *C11orf70* and the *GFP* sequence. The *IFT46* tagged -*GFP* gene was expressed under its own regulators. The *IFT46* (*GSPATG00024708001*) putative promotor was cloned between the *BamHI* and *SphI* sites of pPZZ-GFP02, a modified pPXV-GFP vector (gift of J. Cohen), upstream of the *GFP* sequence. The full length *IFT46* was cloned between *KpnI* and *SmaI* sites of pZZ-GFP02 after the *GFP* sequence. Individual *ND7*-1 mutant cells unable to discharge their trichocysts were then transformed with the expression vectors, by micro-injection into their macronucleus, with filtered and concentrated DNA containing a mixture of the linearized plasmids of interest (5 µg/µl) and of plasmid DNA directing the expression of wild type *ND7*. 3 /4 divisions after the injections, the cell lines issued from the injected cells were screened for their ability to discharge their trichocysts and to express the fluorescent protein. Microinjection was made under an inverted Nikon phase-contrast microscope, using a Narishige micromanipulation device and an Eppendorf air pressure microinjector. Transformants with the same growth rate as untransformed cells and various plasmid DNA copy numbers were selected for experiments. To deciliate the paramecia, they were placed in 10mM Tris pH 7.4, 1mM CaCl2, 5% ethanol for 30 seconds with vortexing. To allow ciliary regrowth, cells were incubated in growth medium for various times from 15 to 30 minutes. Ciliary growth was stopped by fixation and the cells processed for IF. For biochemistry, paramecia were deciliated according to Adoutte et al 1980.^61^ Cilia were recovered by centrifugation at 28000g for 30 minutes, either in SDS-Laemmli buffer or extracted in 20mM Tris pH7.5, 1mM EDTA and 0.2% Triton X-100. Insoluble proteins were precipitated using 9 volumes of methanol before resuspension in SDS-Laemmli buffer. Insoluble proteins were directly solubilized in the same volume of SDS-Laemmli buffer.

### Functional Characterization of C11orf70 in *Chlamydomonas*

Genomic sequences for *Chlamydomonas C11orf70* ortholog FBB5 (Cre12.g556300) and intron/exon boundaries were based on genome assembly annotation version 5.5 and obtained from the Phytozome server. Alignments with FBB5 protein (XP_001694050) orthologs were generated with MEGA version 6.^62^ The expression vector pDIC2linkerHA was created by replacing the *Chlamydomonas* PsaD promoter and 5’UTR in pGenDlinkerHA ^18^ with the promoter and 5’ UTR of the *Chlamydomonas* ODA-IC2 gene.^63^ The resulting expression vector retains a unique Eco RI site between the IC2 ATG start codon and sequences that encode three copies of an HA epitope tag. Two overlapping gBlocks Gene Fragments (Integrated DNA Technologies, Coralville, IA) of 464 and 458 bp were used to insert FBB5 coding sequences at this Eco RI site. The resulting FBB5-HA minigene retains the first intron of the endogenous FBB5 gene. In the resulting protein, the first three amino acids of native FBB5 are replaced with vector-encoded amino acids (MTA -> MVG) and the C-terminal residue (W) is followed immediately by a 9 amino acid linker (AFPRGGISR) and the three HA tag sequences. This expression construct, pFBB5HA, was co-transformed with ARG plasmid pJD67 into an arg2 mutant *Chlamydomonas* strain. Of 72 randomly selected ARG colonies, 8 expressed an HA tagged protein of the expected size (35 kDa), as seen by blots of whole cell samples probed with rat anti-HA McAb 3F10 (Roche, Indianapolis, IN), and one strain was selected for all subsequent work. Mutant strains expressing FBB5HA were generated by standard genetic crosses, using blots of whole cell samples to select tetrad products that expressed the FBB5HA transgene.

*Chlamydomonas* flagellar isolation by dibucaine deflagellation and flagellar fractionation by detergent, followed methods previously described by Mitchell et al., 2005.^64^ *Chlamydomonas* strains *ida1, oda3, oda6, oda8, oda10, oda16* and *fla10*^*ts*^ are available from the *Chlamydomonas* Resource Centre, University of Minnesota. An *ift46* mutant strain expressing an N-terminally truncated IFT46 transgene, and a Guinea pig antibody that recognizes the IFT46 C-terminus were generously provided by George Witman, UMASS Medical School, Worcester, MA.^65^ To test the effects of loss of IFT proteins from flagella, wild type and temperature-sensitive IFT kinesin mutant *fla10*^*ts*^ strains expressing FBB5HA were incubated at 31° C for 2 hr prior to flagellar isolation. Anti-IC2 mouse McAb C11.4^66^ and anti-ODA16 rabbit sera^39^ have been previously characterized.

## Results

### Targeted next generation sequencing identifies *C11orf70* mutations in PCD patients

A targeted next-generation sequencing gene panel was used for mutational analysis in a cohort of affected individuals with PCD. As previously described, the panel contains all the currently known PCD genes and a set of other novel cilia motility disease candidate genes predicted to play a role in cilia motility, based upon collaborative information derived from past patient screening projects and a number of ciliate organism genome and proteome projects.^30^ Screening of 161 unrelated individuals with a confirmed or presumed diagnosis of PCD revealed variants in a new candidate gene *C11orf70* (NM_032930, NP_116319) in two unrelated cases where PCD diagnosis had been confirmed by a number of diagnostic clinical testing modalities.^41^

Firstly, we identified a homozygous *C11orf70* missense variant c.776A>G; p.His259Arg in one of two affected siblings in a Pakistani consanguineous family where the parents are first cousins. Sanger sequencing confirmed the homozygous status of this mutation in the patient and in their affected sibling as well as the carrier status of both parents and an unaffected sibling (**Figure 1A**). This variant is found only once in the ExAC control exome database in a heterozygous carrier from the South Asian population, showing an overall allele frequency in unaffected controls of 8.242e-06. It is absent from the EVS, 1000G and dbSNP control databases. Phylogenetic analysis revealed that the His259 amino acid is a highly conserved residue, located within the DUF4498 domain of *C11orf70* (**Figure 1B**). Using Polyphen-2 software, this mutation is predicted to be damaging, with a score of 0.997 (sensitivity: 0.27; specificity: 0.98). It is also predicted to have a damaging effect on the protein using the SIFT tool. Mutationtaster predicts it to be a disease-causing mutation and it has a CADD score of 23.9 (CADD score of ≥20 indicates a variant is amongst the top 1% of deleterious variants in the genome).

**Figure 1.**
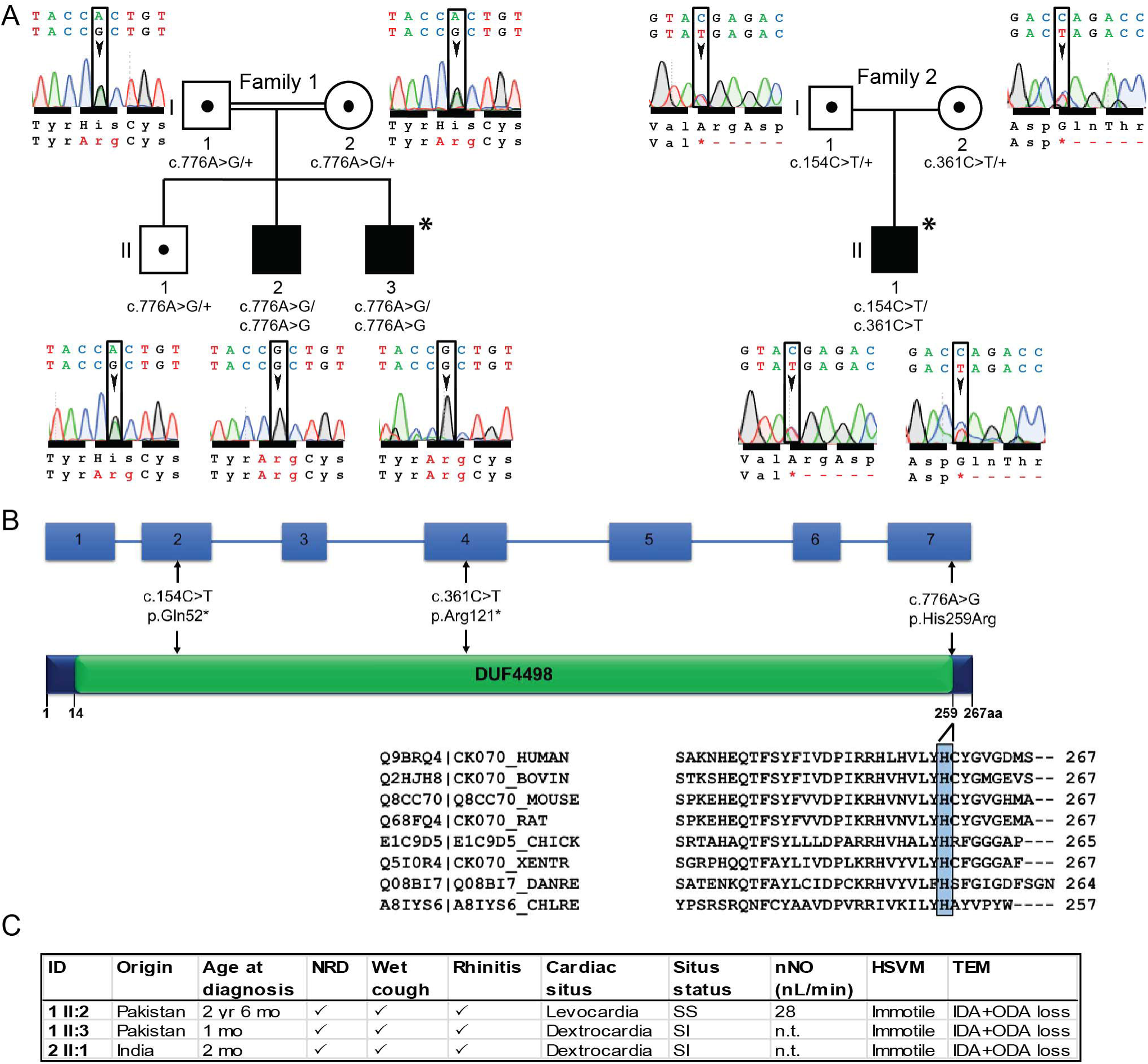
Bi-allelic *C11orf70* mutations identified in individuals with primary ciliary dyskinesia. (A) Pedigree structures of affected family 1 and 2. Electropherograms show variant regions in 1:I.1, 1:I.2 (carrier father and mother), 1:II.1 (carrier sibling), 1:II.2 and 1:I.3 (affected siblings), 2:I.1, 2:I.2 (carrier father and mother), 2:II.1 (affected). Familial segregation shows recessive inheritance in the affected siblings in family 1 (II.2 and II.3) both of whom carry a homozygous missense mutation c.776A>G (p.His259Arg) and in the affected child in family 2 (II.1) who carries compound heterozygous nonsense mutations c.154C>T (p.Gln52*) and c.361C>T (p.Arg121*). (B) The genomic structure of *C11orf70* and the C11orf70 protein shows the locations of the identified disease variants. The only known conserved domain is a ‘domain of unknown function’ (DUF4498). The phylogenetic conservation of the mutated amino acid histidine 259 across different mammalian and other ciliate species is highlighted in blue below the protein. XENTR, *Xenopus tropicalis*; DANRE, *Danio rerio*; CHLRE. *Chlamydomonas reinhardtii*. (C) Clinical characteristics of individuals carrying biallelic *C11orf70* mutations. NRD, neonatal respiratory distress; SS, situs solitus; SI, situs inversus totalis; nNO, nasal nitric oxide level (normal cutoff is >77 nL/min); HSVM, high-speed video microscopy; n.t., not tested as subject too young; IDA, inner dynein arm; ODA, outer dynein arm.

Secondly, we identified compound heterozygous *C11orf70* stop-gain (nonsense) variants in an affected child from an Indian non-consanguineous family, c.154C>T; p.Gln52* inherited from the father and c.361C>T; p.Arg121* inherited from the mother (**Figure 1A**). Sanger sequencing confirmed the mutations in the patient’s DNA and the carrier status of the parents. In the ExAC database, the p.Gln52* mutation (rs767760877) is found in 2 heterozygote carriers from the South Asian population with a total allele frequency of 1.653e-05 and the p.Arg121* mutation (rs561237622) is found only once in a heterozygote carrier from the South Asian population with a total allele frequency of 8.254e-06. Both stop-gain variants are also located within the *C11orf70* DUF4498 domain (**Figure 1B**) and are expected to cause a major deleterious effect on the protein. Their respective CADD scores are 36 (c.154C>T; p.Gln52*) and 39 (c.361C>T; p.Arg121*).

### *C11orf70* mutations cause PCD with combined loss of outer and inner dynein arms

The affected individuals from both families presented with typical PCD symptoms from early life, including a positive history of neonatal respiratory distress and displayed other disease features of chronic wet cough and rhinitis (**Figure 1C**). Only the eldest affected child in family 1 was of age appropriate for measurement of nasal nitric oxide levels which were highly reduced (below the diagnostic cut off of 77 nL/min).^67^ Dextrocardia or situs inversus were observed in both the affected families. High speed video microscopy (HSVM) examination of nasal brushing biopsies of the patients showed that the respiratory epithelium cells had completely immotile cilia (**Movies S1, S2, S3**) compared to the effective ciliary beating observed in both the unaffected sibling 1:II.3 from family 1 (**Movie S4**) and controls (**Movie S5**).

Ultrastructural studies using transmission electron microscopy (TEM) analysis of nasal brushing biopsies, with quantification of outer and inner dynein arms, was performed within a formal clinical diagnostic setting, typically evaluating 50-70 cilia per individual. The 9+2 microtubule pattern of the cilia appeared to be undisturbed overall but the survey revealed a significant loss of both the inner and outer dynein arm structures from respiratory epithelial cell cilia that affected both families equivalently. Outer dynein arms were absent in 67% and 69% of cross sections in family 1 individuals 1:II.2 and 1:II.3 and in 94% of cross section in the family 2 case 2:II.1. The inner dynein arms were lost in 75% and 95% of cilia cross sections in 1:II.2 and 1:II.3 and in 89% of cilia cross sections in 2:II.1. These data are shown in **Figure S1,** with representative images shown in **Figure 2A**.

**Figure 2.**
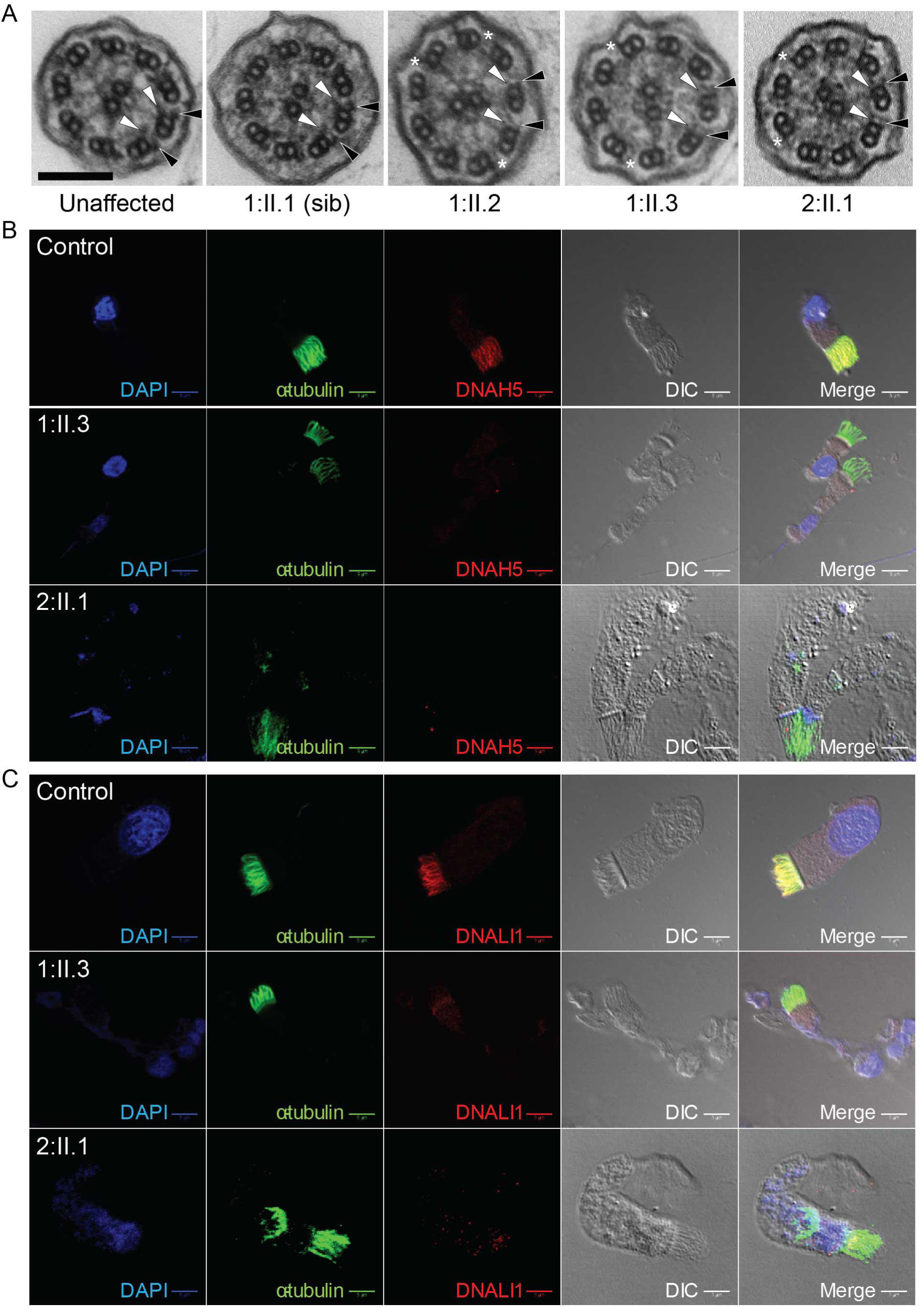
Affected individuals with *C11orf70* mutations display loss of the cilia outer and inner dynein arms in respiratory cells. (A) Representative images of transmission electron micrographs of respiratory cilia in cross section reveals a normal 9+2 pattern of the axoneme but with a loss of both the outer (black arrowheads) and inner (grey arrowheads) dynein arms from the affected patients in family 1 and 2 (right three panels), compared to a healthy control (far left) and the unaffected carrier sibling in family 1 (second left). Scale bar, 100 nm. (B, C) Immunofluorescence staining of respiratory epithelial cells for an antibody marker of the ciliary axoneme (acetylated alpha-tubulin, green) and the outer dynein arm antibody marker DNAH5 (red) shows highly reduced staining in affected patients from both families indicating a loss of the outer dynein arms in agreement with the TEM data (B). Double antibody staining for acetylated alpha-tubulin (green) and the inner dynein arm antibody marker DNALI1 (red) shows a highly reduced staining in affected patients from both families for DNALI1, indicating also a loss of the inner dynein arms (C). Differential interference contrast (DIC) imaging shows the outline of the cell and cilia in each case and the nuclei are in blue (4',6-diamidino-2-phenylindole stain, DAPI). Scale bars, 5 µM.

The composition of the cilia in the respiratory epithelial cell samples in these affected individuals was also examined by immunofluorescence (IF) analysis compared to controls. Respiratory epithelial cells were double-labelled with antibodies directed against the axoneme (acetylated alpha-tubulin) and markers of the outer dynein and inner dynein arms. This showed a significantly reduced level (absence) of staining for the established diagnostic outer dynein arm marker DNAH5 (**Figure 2B**) and inner dynein arm marker DNALI1 (**Figure 2C**) in the patients’ respiratory cilia compared to control samples from an unaffected healthy individual. In agreement with the TEM ultrastructure results, immunofluorescent staining with markers of other cilia structures including the radial spokes (RSPH4A antibody^68^) and nexin-dynein regulatory complexes (GAS8 antibody^69^), did not display altered distribution in the cilia (data not shown).

### *C11orf70* has a characteristic transcriptional profile during mucociliary differentiation

Previous gene expression profiling studies on human tissues showed higher expression of *C11orf70* in multiciliated tissues compared to other non-ciliated tissues.^70; 71^ We analysed the transcriptional profile of *C11orf70* during human motile ciliogenesis, by quantitative PCR of mRNA isolated from air-liquid interface cultures of human ciliated epithelial cells using qPCR primers listed in **Table S1**. We found that the *C11orf70* has a similar expression profile to *DNAH5* (**Figure 3A**) encoding the DNAH5 ODA heavy chain dynein, which is the most commonly mutated PCD gene.^9; 72^ The expression of both genes increased over the time of culture in parallel, with a sharp rise around day 19 at the point when the cells start to ciliate then fell to a plateau phase around day 24.

**Figure 3.**
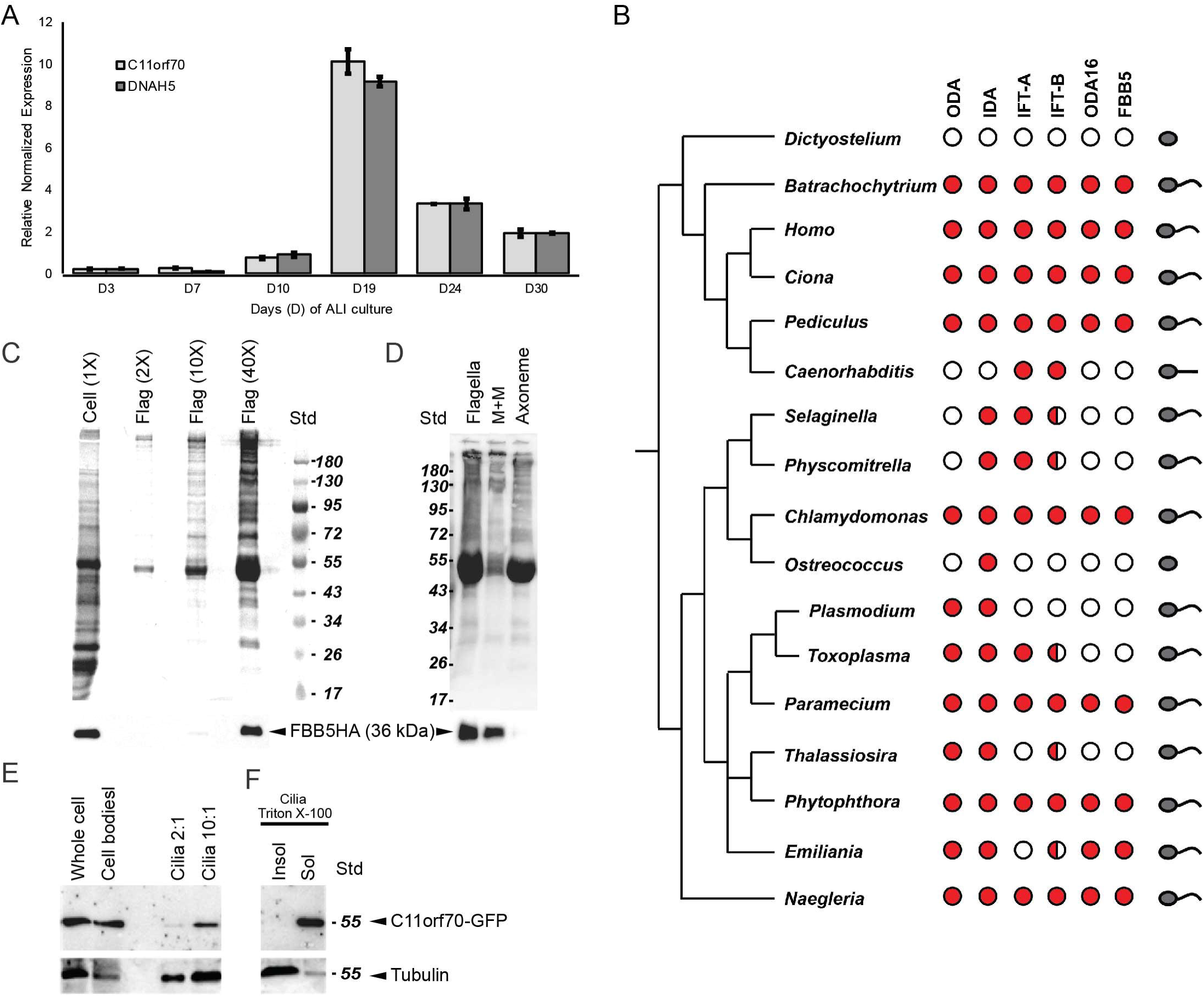
C11orf70 is widely conserved and distributed in both the cytoplasm and cilia. (A) Quantitative RT-PCR analysis of *C11orf70* expression during multi-ciliogenesis in ALI culture over 30 days (D) in culture. The qPCR analysis compared expression of C11orf70 to that of DNAH5 gene. A similar transcriptional profile was seen over time in ALI culture, from around day 10 with a peak at day 19 followed by a plateau from day 24. Error bars indicate SEM. (B) Phylogenomic distribution of *C11orf70* orthologs compared with axonemal dyneins and dynein trafficking proteins. A single *C11orf70* gene copy is retained (filled circles) in genomes of organisms with motile cilia (last column, curved organelle) that use outer dynein arms (ODA column) unless IFT-B complexes are also completely missing (open circles) or compromised by missing subunits (half-filled circles). Each circle represents the results of reciprocal BLAST searches of two or more proteins in each complex, starting with the *Chlamydomonas* sequences (see Methods for details). (C) Stained gels (top) and blot probed for HA (bottom) of cell body and flagellar fractions of *Chlamydomonas* cells expressing tagged *C11orf70* ortholog FBB5-HA. Over 90% of FBB5-HA is cytoplasmic, but a small fraction is present in flagella. (D) Gel (top) and blot (bottom) showing fractionation of isolated *Chlamydomonas* flagella by detergent. FBB5-HA is quantitatively released into the detergent-soluble membrane-matrix fraction. (E) Western blot of deciliated wild type Paramecia (Cell body) expressing C11orf70-GFP as well as cilia fraction of the same cells loaded with a 2-or 10-fold excess of cilia, probed with anti GFP and anti PolyE tubulin antibodies. *C11orf70* is enriched in the cytoplasm, despite 2% is found in the cilia fraction. (F) Cilia fraction in *Paramecium* solubilized using Triton X-100 buffer. The Triton X-100 soluble (Sol) and insoluble (Insol) fractions were loaded at equal stoichiometry. *C11orf70* is found in the soluble fraction corresponding to the matrix and membrane fraction, while PolyE tubulin is highly enriched in the axonemal fraction.

### C11orf70 is highly conserved and expressed in species that require dynein arms for cilia motility

Cross species phylogenetic analysis showed that C11orf70 orthologs are retained in most species that have motile cilia with a few notable exceptions (**Figure 3B and S2**). C11orf70 is not found in the genomes of lower plants that lack outer arms (e.g. *Caenorhabditis, Selaginella, Physcomitrella*) and is also lost from organisms that have outer dynein arms but that do not use IFT for flagellar assembly (e.g. *Plasmodium, Toxoplasma, Thalassiosira*). ODA16, an IFT-interacting protein involved in dynein trafficking, shows a similar distribution. Most of the genomes contained a single C11orf70 copy, with the exception of genomes such as *Xenopus tropicalis* that have undergone duplications.

### C11orf70 localises to both the cilia and cell body in both *Chlamydomonas* and *Paramecium*

Since C11orf70 is a previously uncharacterised human protein, we next proceeded to investigate its wider biological role in cilia motility. Two unicellular model organisms were employed for this analysis, *Chlamydomonas,* a biflagellate alga commonly used to model PCD disease processes^18; 21; 22^ and the ciliate *Paramecium tetraurelia*. *Paramecium* is a unicellular multiciliated organism which is easy to cultivate and, like *Chlamydomonas*, allows for ciliary molecular and biochemical analyses. Moreover, RNAi by feeding in *Paramecium* is a very robust and easy method for functional characterization of potential candidate PCD proteins.^73^

Both the *Chlamydomonas* and *Paramecium* genomes encode a single, well-conserved C11orf70 ortholog. In *Chlamydomonas* it was previously designated as FBB5 based on its phylogenomic distribution with other genes encoding <u>f</u>lagellar and basal body proteins.^74; 75^ *FBB5* is upregulated following deflagellation^76; 77^ but is not present in any published ciliary proteomes. To explore the function of C11orf70 in *Chlamydomonas* and *Paramecium,* we expressed a C-terminal hemagglutinin (HA)-tagged version in wild type *Chlamydomonas* and a C-terminal GFP-tagged version of the *Paramecium* C11orf70 ortholog in the wild type *Paramecium.* Western blots of deflagellated cells and isolated flagella from *Chlamydomonas* revealed that FBB5-HA is predominantly cytoplasmic, with only about 2% appearing in the flagellar fraction (**Figure 3C**). Consistent with these results, GFP-tagged *Paramecium* C11orf70 is present mostly in the cytoplasm with about 2% in cilia (**Figure 3E**). *Chlamydomonas* flagellar FBB5-HA was quantitatively released by detergent treatment, which placed it in the soluble membrane and matrix fraction (**Figure 3D**). Similarly, cilia purification followed by Triton X-100 extraction in *Paramecium* released the ciliary C11orf70 into the soluble fraction (**Figure 3F**).

### *C11orf70* has a conserved role in dynein arm assembly in *Paramecium*

We further assessed the involvement of *C11orf70* as a novel causative gene of PCD and its role in cilia motility by gene silencing in *Paramecium*. RNAi knockdown of the *Paramecium C11orf70* reduced the expression of *C11orf70* by >80% (**Figure 4A**). As a control, the *ND7* gene, which plays no role in cilia motility, was subjected to RNAi in parallel. *C11orf70* knockdown led to an evident phenotypic defect in the *Paramecium* swimming pattern with severely reduced velocity after 72 hr (**Figure 4B**). The average swimming velocity of the *C11orf70*-RNAi cells (0.58±0.22 mm/s) was reduced to 68% of the control *ND7*-RNAi cells (0.85±0.20 mm/s) (**Figure 4C**). This decrease in the *Paramecium* swimming speed after *C11orf70* RNAi was confirmed not to be a consequence of an alteration in cell size, cilia length or number of cilia per cell (**Figure S3**). When HSVM analysis of the cilia motility of *C11orf70* RNAi-silenced *Paramecium* was performed, a significant decrease in cilia beat frequency (CBF) was observed in the *C11orf70* knockdown cells compared to the control *ND7* knockdown cells (**Figure 4D**). The mean CBF of the control ND7-RNAi *Paramecium* was 24.3±2 Hz whilst the cilia of the *C11orf70*-RNAi *Paramecia* were beating at a frequency averaging 20.3±4 Hz. Despite of the modest reduction of the cilia beat frequency, a close examination of the movies showed that some cilia were not beating. The cilia waveform was also affected by *C11orf70* knockdown with a variable phenotype ranging from a slight reduction in amplitude to a half-beat (**Movies S6, S7**).

**Figure 4.**
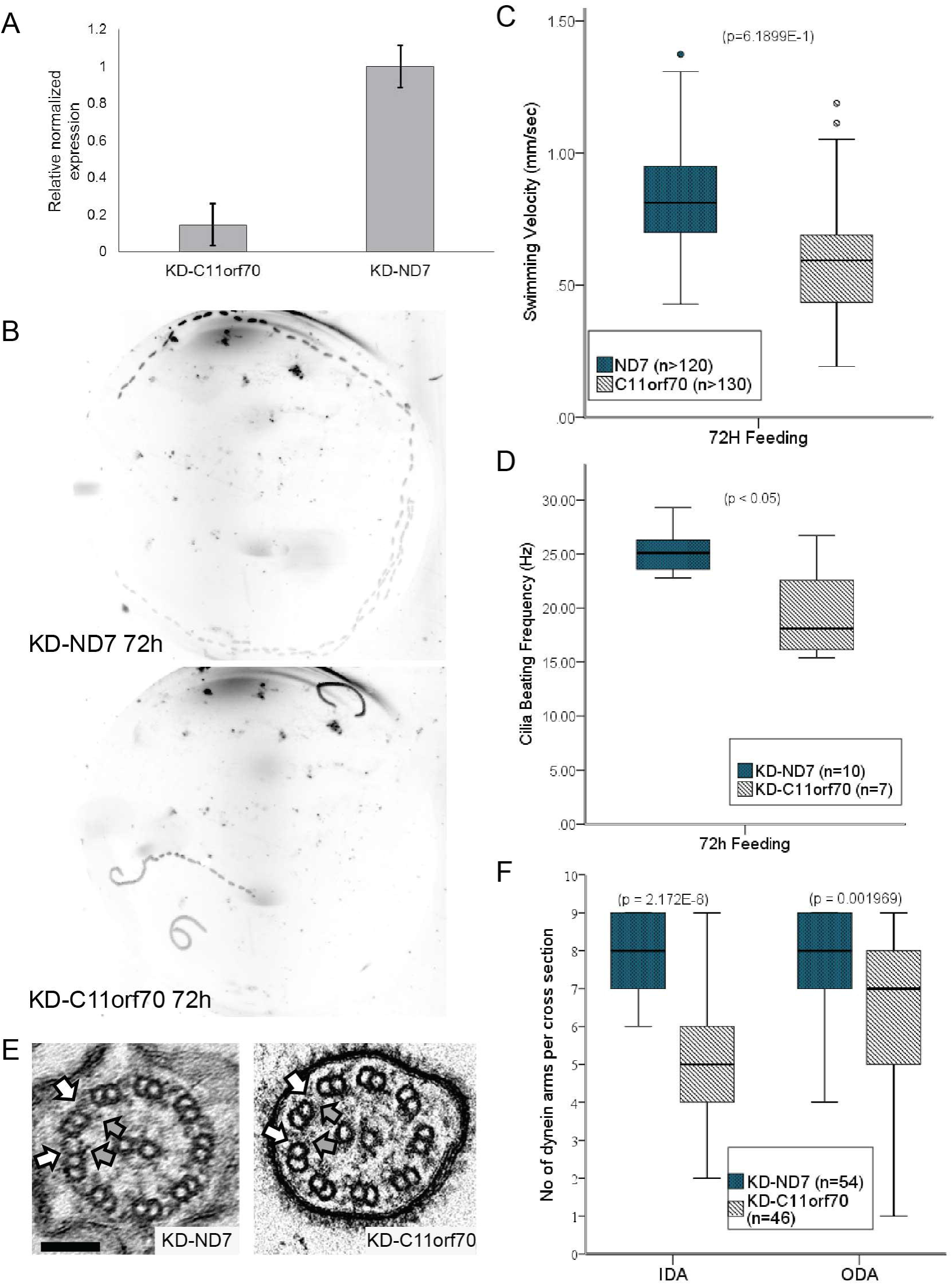
Paramecium gene knockdown shows a conserved function for C11orf70 in dynein arm assembly. (A) Quantitative RT-PCR analysis of *C11orf70* expression in both the *C11orf70* knockdown cells and the *ND7* knockdown control. The expression level of *Paramecium C11orf70* is reduced by about 80% in the *C11orf70* knockdown cells compared to its expression in the control *ND7* knockdown cells. (B) Z projection of track recording Paramecium swimming under a dark-field microscope using a 10X objective after 72 hr of RNAi treatment. *C11orf70* silenced cells (bottom panel) show a severe motility phenotype compared to the control *ND7* silenced cells (top panel). (C) Analysis of the swimming velocity of *C11orf70* and *ND7* knockdown cells. The swimming velocity of *C11orf70* knockdown cells is reduced by 32% of that of the control *ND7* knockdown cells. At least 90 organisms evaluated per condition; P<0.001, Independent samples t test. (D) At 72 hours of RNAi the ciliary beat frequency of *C11orf70* knockdown cells is reduced by about 20% compared to *ND7* control knockdown cells. 7-10 paramecia evaluated per condition; P<0.05, Mann-Whitney U-test. (E) Transmission electron micrographs of *Paramecium* cilia in cross section showing normal 9+2 arrangement for the control *ND7* knockdown (left) and absence of the outer (white arrows) and inner (grey arrows) dynein arms in the *C11orf70* knockdown cells (right). Scale bar, 100 nm. (F) Quantification of TEM dynein arms counts across >40 cross sections per strain showed significant alteration of numbers of ODA and IDA in *C11orf70* knockdown cells compared to ND7 control knockdown cells. P>0.001, Independent samples t test.

These results led us to examine the presence of cilia ODA and IDA by TEM after *C11orf70* knockdown, since ODA are supposed to be involved in the beat frequency and IDA in the waveform.^65; 66^ TEM examination of cilia cross sections showed a similar phenotype to that seen in individuals carrying *C11orf70* mutations (**Figure 4E**), with a statistically significant reduction of both IDA and ODA numbers per cilia cross section observed in the *C11orf70* knockdown cells compared to ND7 knockdown strain. The mean numbers of IDA and ODA were 7.8 ± 1.1 and 7.9 ± 1.3 in the control *Paramecium* cilia cross sections compared to 4.9 ±1.8 and 6.2± 2.2 in the *C11orf70* silenced *Paramecium* cilia cross sections respectively (**Figure 4F**).

Overall, these phenotypic findings in P*aramecium* after *C11orf70* ortholog silencing indicate the high level of evolutionary conservation of its gene function across ciliate species, with cilia defects that are consistent with the diagnostic findings in individuals affected with PCD that carry mutations in *C11orf70*.

### C11orf70 has a conserved intracellular distribution pattern similar to IFT-associated proteins

To gain insight into the mechanism by which C11orf70 affects inner and outer dynein arm assembly, we next studied its behaviour during cilia re-growth in *Paramecium.* We compared its behaviour in comparison to GFP alone, which enters cilia by diffusion, and IFT46-GFP, one member of the IFT-B complex (involved in IFT-associated anterograde transport of ODA into cilia), by expressing these along with a GFP-tagged C11orf70 protein. Several transformant clones showing a wild-type growth phenotype and expressing various levels of C11orf70-GFP were analysed. All of them showed the same results that the tagged C11orf70-GFP protein is found mostly in the cytoplasm with a small amount found in cilia (**Figure 5A**), in agreement with the *Chlamydomonas* and *Paramecium* cell fractionation experiments in **Figure 3B, C and D**. We then assessed the localization of C11orf70 after deciliation followed by cilia re-growth of the transformed *Paramecium* cells. After 15 mins of cilia re-growth, when IFT transport is particularly active, we observed that IFT-46 GFP accumulates at the ciliary tips (**Figure 5B, right panels**) compared to a uniform distribution of fluorescence in the cilia of cells transformed with GFP alone (**Figure 5B, left panels**). Similarly to IFT46-GFP, the C11orf70-GFP protein showed a comparable cilia tip accumulation during ciliogenesis (Figure 5B, middle panels).

**Figure 5.**
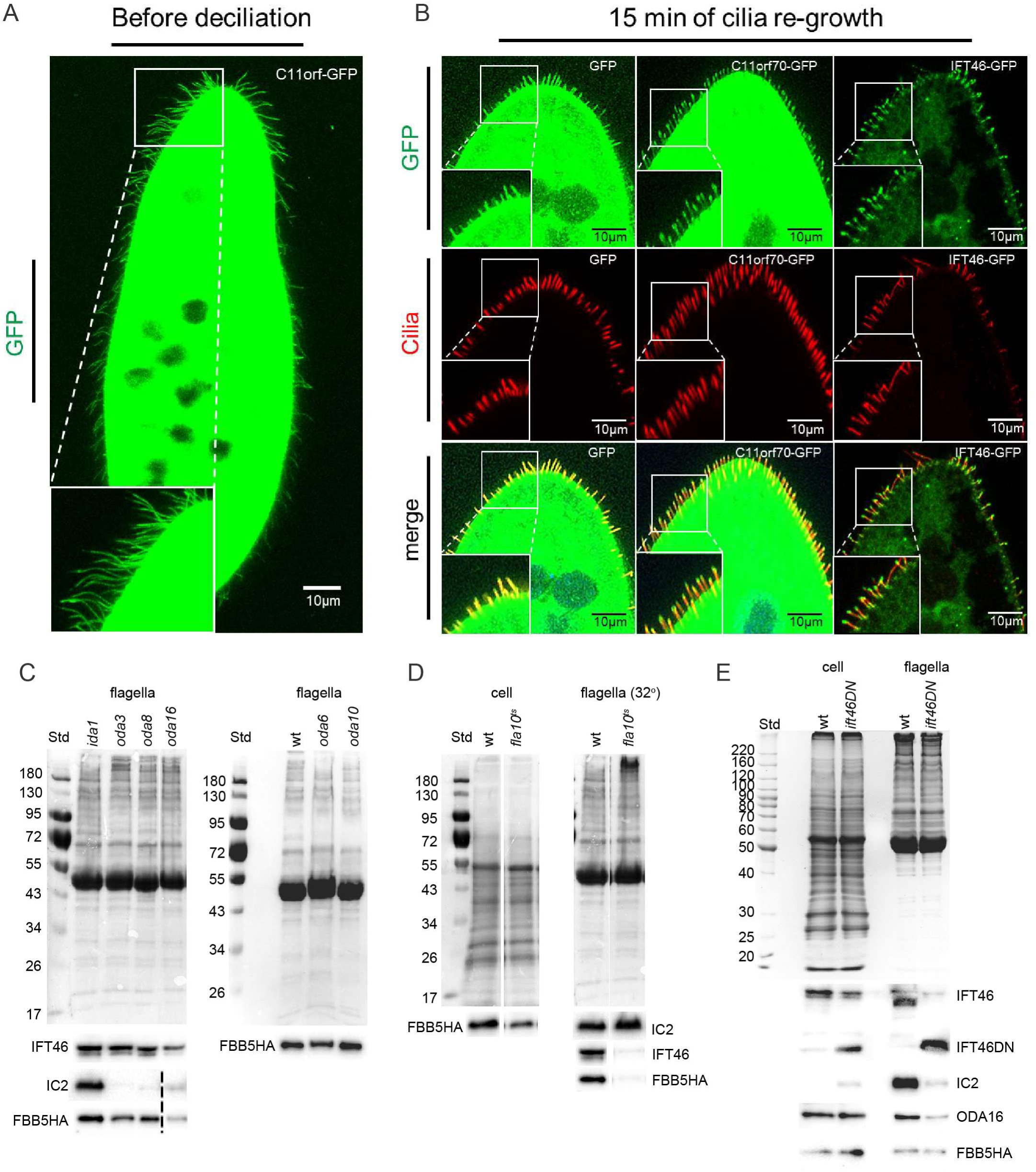
C11orf70-GFP accumulates at the ciliary tip during reciliation and its flagellar abundance is IFT-dependent. (A) Paramecia expressing the *Paramecium* C11orf70-GFP show a very high level of cytoplasmic GFP staining with protein also present in the cilia. The cell body was overexposed in order to allow the detection of cilia. (B) *Paramecium* expressing either GFP alone **(left panels**), C11orf70-GFP (**middle panels**) or IFT46-GFP (**right panels**) were subject to deciliation after which the cilia were allowed to regrow for 15 minutes, in order to study the localization of the tagged proteins during reciliation. Direct GFP fluorescence (green, top panels) and monoclonal antibody staining against polyglutamylated tubulin (red, middle panels) are shown along with the merge (bottom panels). Paramecia expressing GFP are used as a negative control since the GFP protein enters the cilia passively (<27 kDA). The GFP staining is found distributed homogeneously along the entire length of the cilia. In contrast, IFT46-GFP accumulates at the ciliary tips during ciliary regrowth, as expected for an IFT-B family member. C11orf70-GFP is found at much higher levels in the cytoplasm than IFT46-GFP and is also present in the cilia. Upon ciliary regrowth, C11orf70- GFP behaves in a similar fashion to IFT46 GFP in cilia with an accumulation of both proteins observed at the ciliary tips. Scale bars, 10 µm. (C) Coomassie stained gels (above) and western blots (below) show that the flagellar fraction abundance of *Chlamydomonas* FBB5 is similar strains carrying mutations in components of the outer or inner dynein arms (*oda6, ida1*), the ODA-DC (*oda3*) and late-stage ODA assembly factors (*oda8, oda10*). However, its abundance is reduced by 75% in the absence of ODA16, an IFT-associated dynein transport factor (left panel). Blots of IFT46 used as a marker of flagellar IFT, shows that IFT is reduced by 50% in *oda16*. Dotted line indicates removal of an intervening lane on the blot. D) Gels (top) and blots (bottom) of whole cells and flagella from wild type and *fla10*^*ts*^ strains of *Chlamydomonas*. Similar levels of FBB5-HA are expressed in both strains (left panels, whole cells). Flagella from *fla10*^*ts*^ cells incubated at the restrictive temperature lack IFT46 and have also lost FBB5-HA, whereas elevated temperature has no effect on wild type flagella (right panels). (E) Gels (top) and blots (bottom) of whole cells and flagella from FBB5-HA-expressing wild type *Chlamydomonas* and an *ift46* mutant strain that also express an N-terminally truncated IFT46 transgene. Similar levels of IFT46, IFT46∆N, ODA-IC2, ODA16 and FBB5-HA are expressed in whole cells of both strains (left panels). The truncated IFT46 supports IFT-dependent flagellar assembly, but the flagella show reduced levels of ODAs (IC2) and ODA16. FBB5-HA levels, however, are not affected (right panels).

### Flagellar C11orf70 abundance is IFT dependant

We next took advantage of *Chlamydomonas* genetics to investigate the functional role of *C11orf70* in IFT-related cilia motility by expressing the HA-tagged FBB5 ortholog in various axonemal assembly mutant strains. We found that flagellar FBB5-HA levels were not affected by mutations in outer or inner dynein arm subunits (*oda6, ida1* mutant strains)^78; 79^, the ODA-DC (*oda3*)^80^ or late-stage cytoplasmic assembly factors (*oda8, oda10*).^37; 80^ However FBB5-HA levels were found to be decreased by a loss of IFT-associated dynein transport factor ODA16 (*oda16*)^39; 40; 81^ (**Figure 5C**), with confirmation in the *oda16* strain that FBB5-HA was expressed at equivalent levels to the other tested strains (**Figure S4**). Since this suggests a role for FBB5 in IFT-dependent dynein transport, we further tested the role of IFT in transport of FBB5 to the flagellar matrix by expressing FBB5-HA in the temperature-sensitive IFT kinesin mutant deficient in anterograde IFT, *fla10*^*ts*^.^82^ After shifting to the restrictive temperature (32º) for 1.5 hr, the *fla10*^*ts*^ flagella were found to be depleted of IFT complexes (monitored by the abundance of IFT46) and of FBB5-HA (**Figure 5D**). Since ODA transport factor ODA16 interacts with *IFT46*^39^ and specifically depends on an interaction with the N-terminal domain of IFT46 for ODA transport, ^65; 81^ we tested the effects of deleting the IFT46 N-terminus on flagellar abundance of FBB5. Flagella of an *ift46* strain that also expressed an N-terminally truncated IFT46 protein^65^ had reduced levels of ODA16 and greatly reduced assembly of ODAs (as monitored by the abundance of ODA intermediate chain IC2), but showed no reduction in FBB5-HA (**Figure 5E**).

## Discussion

Here we report mutations in a previously uncharacterised gene, *C11orf70*, in individuals with PCD and situs inversus, demonstrating the potential power of using a targeted NGS approach for novel PCD gene discovery. *C11orf70* mutations give rise to immotile respiratory cilia that have a normal 9+2 ultrastructure with a combined lack of both inner and outer dynein arms and affected individuals have immotile cilia.

Cilia axonemal structure is highly conserved across species and various model organisms have been used in the past to characterize candidate genes for PCD. Phylogenetic analysis showed that C11orf70 is a highly conserved protein across anciently diverging species, being present today only in organisms with motile cilia/flagella. It is lost in organisms that do not use IFT assembly of their cilia/flagella. It has a specific distribution across different species, following a similar pattern to that of IFT-associated dynein assembly factors like *ODA16* (*WDR69*).^39; 40; 65^ Together these findings support the importance of C11orf70 in the cytoplasmic assembly, maturation and IFT-B-based transport into cilia of dynein arm motors that are required for cilia motility in multiciliated cells.

Transcriptional profiling of *C11orf70* shows that it is more enriched in tissues with motile cilia than non-ciliated tissues,^70; 71^ although it was not identified in proteomic profiling of human respiratory cilia.^83^ Microarray-based gene expression profiling has shown that *C11orf70* expression is upregulated during mucociliary differentiation.^84^ This is consistent with our finding that the *C11orf70* expression profile follows a similar pattern to that of other PCD genes during ciliogenesis, with a peak around the time of cilia emergence that reaches a plateau afterwards. Interestingly, the *Chlamydomonas C11orf70* ortholog, *FBB5*, has been identified as amongst the flagellar-basal body group of genes, ^74; 75^ upregulated following deflagellation, ^76; 77^ but it is not present in any ciliary proteomes. The *Paramecium C11orf70* ortholog, *GSPATG00011350001*, was also not detected within the cilium proteome^85^ or during cilia regrowth after deciliation of *Paramecium*.^86^

To study the role of *C11orf70* in cilia motility, we knocked down its ortholog in a new ciliate model, *Paramecium*. We first confirmed the efficiency of silencing at the RNA level and then studied the phenotypic consequences of gene silencing on cilia motility and ultrastructure. We found that *C11orf70* RNAi knockdown led to a reduction in swimming velocity as a secondary effect of a decrease in cilia beating frequency. Analysis of the ciliary ultrastructural phenotype demonstrated the highly conserved, ancient function of C11orf70 in dynein arm assembly since *C11orf70*-silenced *Paramecium* cells mimic the combined outer and inner dynein arm loss of affected individuals with PCD carrying *C11orf70* mutations, consistent with their immotile respiratory cilia. This provides evidence that C11orf70 plays a role in the dynein arm assembly/transport process. This work highlights the utility of *Paramecium* for studies of cilia motility and its emergence as a new organism to model the cell biology of disease mechanisms underlying PCD.

In *Chlamydomonas* we found that the distribution of the C11orf70 ortholog FBB5 between the cytoplasm and cilia/flagella was similar to other IFT-associated dynein assembly proteins such as ODA16^39^ and ODA8.^37^ A similar distribution was seen for *Paramecium* C11orf70. Our expression studies in both *Chlamydomonas* and *Paramecium* showed that C11orf70 is a soluble, mainly cytoplasmic protein, with a small amount also present in the matrix fraction of cilia/flagella. The protein was around 40X more abundant in the cytoplasm than the cilia/flagella in both organisms. This is similar to ODA16, which is concentrated around the basal bodies with ~2% found in flagella.^39; 40^ The anterograde IFT partner of ODA16, IFT-B family member IFT46, shows a similar pattern, also concentrated around basal bodies with ~5% present in flagella. ODA16 and IFT46 are known to interact together to mediate the transport of outer dynein arm into the cilia.^65^ Amongst the other PCD-associated dynein assembly factors, the distribution of C11orf70 is reminiscent of two, ZMYND10 and C21orf59, that are considered predominantly cytoplasmic with a small component potentially also detectable within cilia.^32; 34^

The flagellar localisation of *Chlamydomonas* FBB5-HA was unaffected by most mutations that disrupt ODA assembly but was reduced by loss of the IFT46-associated transport factor ODA16, although we found that its abundance in the axoneme was not dependent on the N-terminus of IFT46. We note that this effect could result from a general reduced abundance of IFT in *oda16* flagella rather than being a direct consequence of ODA16 mutation. However in *Paramecium*, we showed that C11orf70 protein accumulates in the cilia tips in the same way as IFT46, suggesting that C11orf70 undergoes active transport to enter the cilia. The IFT-B family including IFT46 governs anterograde (base to tip) transport in the cilium and is very active during ciliation, in order to build the cilium. IFT46-GFP is released at the tip of the cilium where it accumulates during active IFT and C11orf70 may behave similarly.

Together, these data identify mutations in *C11orf70* as a cause of PCD with combined outer and inner dynein loss, cilia immotility and situs inversus. This has important clinical implications for improved understanding of the genetics basis of disease and counselling of affected families. The combined evidence from human and ancient unicellular ciliate organisms support a role for C11rof70 as a new dynein assembly/transport factor, its abundance in the axoneme dependent upon IFT.

## Description of Supplemental Data

1 Supplementary data file 7 Supplementary movies

## Conflicts of Interest

The authors report no conflicts of interest. The authors alone are responsible for the content and writing of the paper.

### Acknowledgements

We are very grateful to the families who participated in this study and thank the UK PCD Family Support Group for their continued support. We thank Ms Dani Lee and Prof Chris O’Callaghan (UCL Great Ormond Street Institute of Child Health) for help with assessment of *Paramecium* motility and Dr Dale Moulding (GOS-ICH Confocal Microscopy Core Facility) for experimental assistance. Funding for this study was provided by Action Medical Research (GN2101; H.M.M.) and Great Ormond Street Children’s Charity grant (V4515; H.M.M.) and Leadership awards (V1299, V2217; H.M.M.). We acknowledge support from the NIHR Biomedical Research Centre at Great Ormond Street Hospital for Children NHS Foundation Trust and University College London (Doctoral Trainee Support Award; M.R.F.). M.R.F. is supported by the British Council Newton-Mosharafa Fund and the Ministry of Higher Education in Egypt. Work by A.S. is independent research funded by a postdoctoral research fellowship from the National Institute of Health Research and Health Education England, and the views expressed in this publication are those of the authors and not necessarily those of the NHS, the National Institute of Health Research or the Department of Health. We would like to thank C. Mathon for generating plasmid constructs, L. Shi for the IFT46 construct, J. Bonaventure for his help in RNA isolation and C. Janke for his generous gift of PolyE antibodies. The authors acknowledge the contribution of the BEAT-PCD COST Action. The authors participate in the COST Action BEAT-PCD: Better Evidence to Advance Therapeutic options for PCD network (BM1407) and this work was supported by two BM1407 COST Action STSM Grants awarded to M.R.F. and P.L.B.

### Web Resources

CILDB: http://cildb.cgm.cnrs-gif.fr/
ParameciumDB: http://paramecium.cgm.cnrs-gif.fr/
RNAi off-target tool: http://paramecium.cgm.cnrs-gif.fr/cgi/tool/alignment/off_target.cgi
dbSNP build 141: https://www.ncbi.nlm.nih.gov/projects/SNP/
ExAC: http://exac.broadinstitute.org/
Exome Variant Server: http://evs.gs.washington.edu/EVS/
1000Genomes: http://1000genomes.org/
NCBI Primer-BLAST tool: https://www.ncbi.nlm.nih.gov/tools/primer-blast/
Human Splicing Finder: http://www.umd.be/HSF3/
SIFT: http://sift.jcvi.org/
Polyphen-2: http://genetics.bwh.harvard.edu/pph2/
Mutation Taster: http://www.mutationtaster.org/
CADD: http://cadd.gs.washington.edu/
MAPP: http://mendel.stanford.edu/sidowlab/downloads/MAPP/index.html
PhastCons: http://compgen.cshl.edu/phast/
PhyloP: http://compgen.bscb.cornell.edu/phast/help-pages/phyloP.txt
Phytozome: https://phytozome.jgi.doe.gov
BLAST/BLOSUM62: https://blast.ncbi.nlm.nih.gov/Blast.cgi
OMIM: http://www.omim.org/
RefSeq: http://www.ncbi.nlm.nih.gov/RefSeq
Ensembl Genome Browser: http://www.ensembl.org/index.html
*Chlamydomonas* Resource Centre: http://chlamycollection.org/
NCBI: https://www.ncbi.nlm.nih.gov/

## Accession numbers

The NCBI accession numbers used in this paper are: NM_032930 and NP_116319 (human C11orf70); XM_001443078.1 and XM_001443078.1 (*Paramecium* GSPATG00011350001); XM_001693998.1 and XP_001694050 (*Chlamydomonas* FBB5).

